# Antimicrobial emulsifier – glycerol monolaurate impacts gut micobiome inducing distinct effects on metabolic syndrome in low-fat diet fed mice

**DOI:** 10.1101/2020.09.11.294454

**Authors:** Zengliang Jiang, Congmei Xiao, Xi Zhang, Minjie Zhao, Tao Liu, Yisong Xu, Hui Zhang, Ju-Sheng Zheng, Fengqin Feng

## Abstract

Previous study demonstrated 150 mg·kg^−1^ glycerol monolaurate (GML) consumption induced gut microbiota dysbiosis and metabolic syndrome (MetS) in low-fat diet fed mice. However, little is known about the dose-effect of dietary GML modulating the gut microbiome alterations and its impacts on the induction of MetS in low-fat diet. Dietary GML-induced effects on MetS and gut microbiota alterations were investigated, combined with antibiotics-treated germ-free experiment and specific gut microbiota transplantation. Our results demonstrated that high-dose (500 mg·kg^−1^) GML alleviated MetS by significantly decreasing the body weight, weight gain, food intake, fat droplet size and percentage of abdominal fat, serum triglycerides (TG), LDL, LPS, TNF-α and atherogenic index, compared to the medium-dose (150 mg·kg^−1^) GML. Importantly, high-dose GML significantly increased *Lactobacillus reuteri* compared to the medium-dose GML. Co-occurrence network analysis revealed *Lactobacillus reuteri* was positively associated with the metabolic improvement of high-dose GML. Notably, antibiotics-treated germ-free experiment and *Lactobacillus reuteri* transplantation demonstrated that altered gut microbiota was necessary and sufficient for GML-induced distinct effects on metabolic syndrome. Our results indicate that GML impacts gut microbiome inducing distinct effects on metabolic syndrome, thereby calling for reassessing the safe dosage of GML and other non-specific antibacterial food additives.

**IMPORTANCE:** Growing evidence indicate that the broad use of food emulsifying agents may lead to increase the societal incidence of obesity/ MetS and other chronic inflammatory diseases. GML is widely and regularly consumed as a generally safe food emulsifier and as a potent antimicrobial agent in commonly foods such as meat products, cereals and soft beverage by the general public. Our results indicate that GML impacts gut microbiome inducing distinct effects on metabolic syndrome. Our study provides important and timely evidence supporting the emerging concept that non-specific antibacterial food additives have two-sided effect on gut microbiota contributing to the uncertainties for the incidence of obesity/metabolic syndrome and other chronic inflammatory diseases.

---

Metabolic syndrome (MetS) is associated with an increased risk of type 2 diabetes and cardiovascular disease (CVD) (1, 2). In 2001, the National Cholesterol Education Program–Adult Treatment Panel III (NCEP–ATP III) proposed that MetS is identified by meeting at least three of the five following metabolic alterations: elevated abdominal obesity, high circulating levels of triacylglycerols, reduced high-density lipoprotein (HDL)-cholesterol, elevated fasting plasma glucose and elevated blood pressure (3). In the United States, nearly 35% of all adults and 50% of those aged 60 years or older were estimated to have the MetS (4). With its prevalence increasing exponentially in recent decades, MetS has been regarded as an important public health issue in the 21st century, affecting about 25% of the population in developed countries (5). Therefore, there is an urgent need to investigate the etiology of the emerging global epidemic.

Noticeably, growing evidence indicates that the broad use of emulsifying agents may lead to increase the societal incidence of obesity/ MetS and other chronic inflammatory diseases (6–8). Glycerol monolaurate (GML), a naturally occurring glycerol monoester of lauric acid enriched in coconut oil, is widely and regularly consumed as a generally safe food emulsifier and as a potent antimicrobial agent in commonly foods such as meat products, cereals and soft beverage by the general public (9, 10). The dosage of GML in food products is not limited and its additive amount in food and health care products ranges from 10 mg·kg^−1^ to 2000 mg·kg^−1^ (21 CFR GRAS 182.4505). Our previous study demonstrated that 150 mg kg^−1^ GML consumption induced MetS in low-fat diet fed mice (11) and 450 mg kg^−1^ GML consumption ameliorated MetS in a high-fat diet fed mice (12). However, it remained unclear the dose-effect relationship of dietary GML on metabolic syndrome in the low-fat diet.

Recent studies revealed that the development of MetS, obesity and metabolic comorbidities were associated with disturbance of the gut microbiota possibly due to the increased energy harvest, production of toxic bacterial metabolites and greater intestinal permeability (13–15). The decrease in beneficial bacteria and increase in pro-inflammatory/pathogenic bacteria leaded to host metabolic disorder and a low grade systemic inflammatory state (16, 17). Interestingly, our previous study demonstrated that a relatively low-dose GML consumption induced gut microbiota dysbiosis, MetS and systemic low-grade inflammation in low-fat diet fed mice (11). Thus, a better understanding of the effect of dietary GML on modulating gut microbiota would be important for evaluating the safety of its use in consumed foods. However, it is still unclear the regularity of dietary GML mediated the gut microbiota alterations and whether GML-mediated gut microbiota alterations were necessary for the induction of MetS.

To directly answer above questions, we studied the GML-mediated modulation of community structure, composition, functional genes of gut microbiota, and the resultant effects on host MetS in mice. Further, the antibiotics-treated germ-free experiment and specific gut microbiota transplantation experiment were carried to prove the necessity of gut microbiota for GML-induced MetS.

## RESULTS

### Dietary GML induces distinct effects on metabolic syndrome

In order to investigate the dose-effect relationship of dietary GML on metabolic syndrome, relatively low (50 mg·kg−1, T1 group), median (150 mg·kg−1, T2 group) and high (500 mg·kg−1, T3 group) doses of GML were utilized to feed mice for 8 weeks. Markedly, the body weight (*p* < 0.001 and *p* < 0.001), weight gain (*p* < 0.001 and *p* = 0.003), food intake (*p* = 0.037 and *p* = 0.047) and abdominal fat (*p* = 0.016 and *p* = 0.032) in the medium-dose group were significantly increased compared to those of the CON, low-dose and high-dose groups, while no significant differences were observed among the CON, low-dose and high-dose groups (Fig. 1a-c,e). Moreover, body fat percentage in the low-dose and medium groups (*p* = 0.013 and *p* = 0.037, respectively), but not in the high-dose group, were significantly increased compared to the CON group (Fig. 1d).

**Figure 1.**
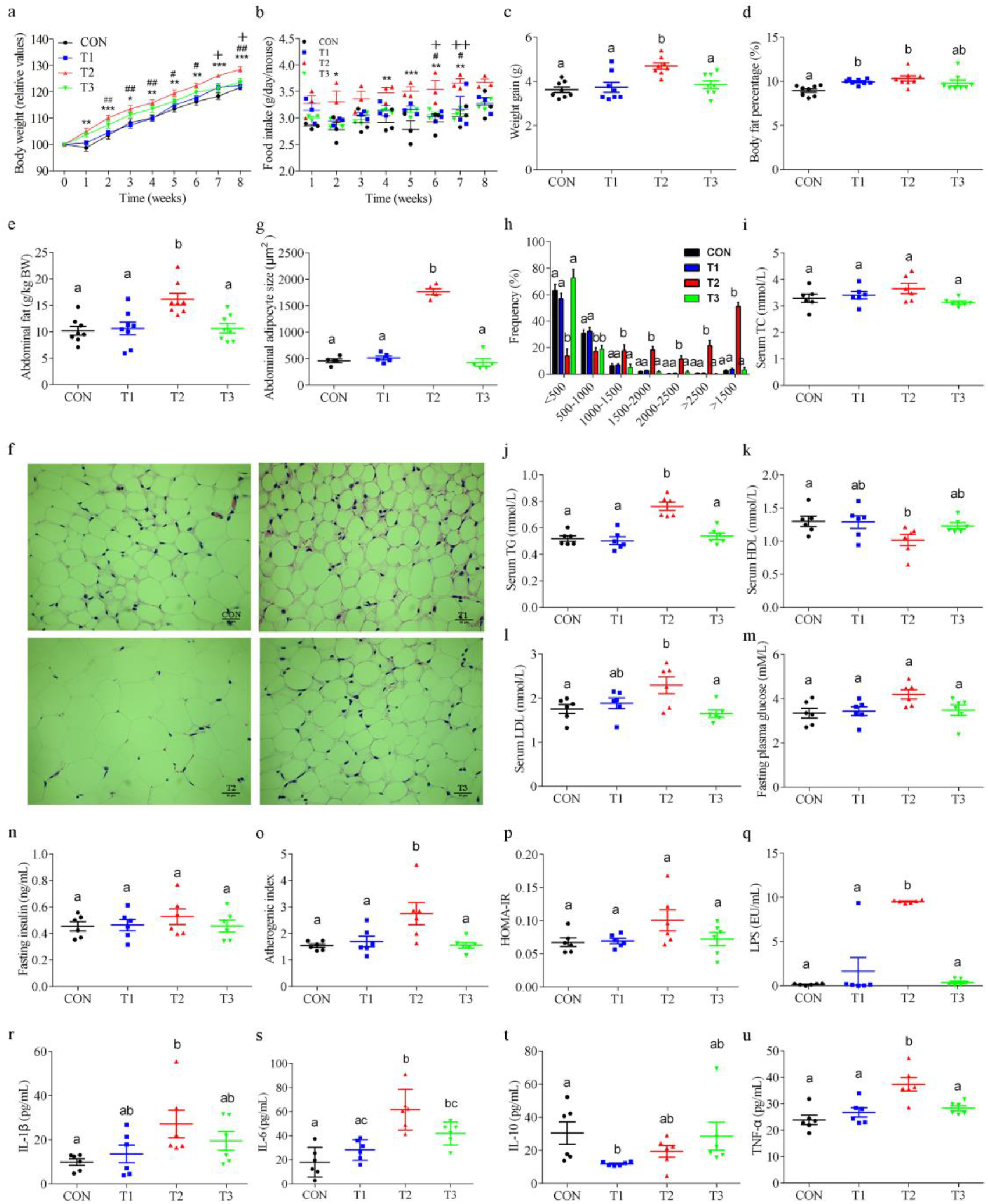
Dietary GML induces distinct effects on metabolic syndrome. Dose-effects of dietary GML (n = 8/group) on **(a)** body weight, **(b)** food intake (n = 3, 3 cages/group), **(c)** weight gain, **(d)** body fat percentage, **(e)** abdominal fat percentage, **(f)** abdominal fat content assessed using H&E staining (n = 5/group; scale bar, 30 μm), the size **(g)** and frequency **(h)** of stained fat droplets in abdominal adipose tissue (n = 5/group), **(i)** serum total serum cholesterol (TC), **(j)** triglycerides (TG), **(k)** high-density lipoprotein cholesterol (HDL-C), **(l)** low-density lipoprotein cholesterol (LDL-C), **(m)** fasting plasma glucose, **(n)** fasting insulin, **(o)** atherogenic index and **(p)** homeostasis model assessment-insulin resistance index (HOMA-IR), serum LPS concentration **(q)**, IL-1β **(r)**, IL-6 **(s)**, IL-10 **(t)** and TNF-α **(u)** (n = 6/group). Data are expressed as mean ± SE. Value with asterisk or different alphabet is significantly different (T2 *VS* CON: \**p* < 0.05, \*\**p* < 0.01, \*\*\**p* < 0.001; T2 *VS* T1: ^#^*p* < 0.05, ^##^*p* < 0.01, ^###^*p* < 0.001; T3 VS T2: ^+^p < 0.05, ^++^p < 0.01, ^+++^p < 0.001).

Markedly, H&E histopathology demonstrated that the size of stained fat droplets of abdominal adipose tissue in the medium-dose group were significantly larger (*p* < 0.001) than that of the CON, low-dose, and high-dose groups (Fig. 1f and g). Meanwhile, the frequency of < 500 μm^2^ of stained fat droplets of abdominal adipose tissue in the medium-dose group were significantly decreased (*p* < 0.001, *p* < 0.001 and *p* < 0.001, respectively), and the frequency of more than 1000 μm^2^ of fat droplets in the medium-dose group were significantly increased, compared to those of the CON, low-dose and high-dose groups (*p* < 0.001, *p* < 0.001, *p* < 0.001, respectively), while no significant differences among the CON, low-dose and high-dose groups were observed (Fig. 1h).

Importantly, the serum TG (*p* < 0.001, *p* < 0.001 and *p* < 0.001, respectively) and atherogenic index (*p* = 0.017, *p* = 0.047 and *p* = 0.023, respectively) in the medium-dose group were significantly higher than those in the CON, low-dose and high-dose groups (Fig. 1j and o). In addition, the serum LDL (*p* = 0.033 and *p* = 0.009) in the medium-dose group were significantly higher than those in the CON and high-dose groups (Fig. 1l) and the serum HDL (*p* = 0.034) in the medium-dose group was significantly lower than that in the CON group (Fig. 1k). However, no significant differences for the serum TC, TG, HDL, LDL fasting plasma glucose, insulin, atherogenic index and HOMA-IR among the CON, low-dose and high-dose groups were observed respectively.

Notably, significant increases (*p* < 0.001, *p* < 0.001 and *p* < 0.001, respectively) were observed for the serum LPS concentration in the medium-dose group compared to the CON, low-dose and high-dose groups (Fig. 1q). Accordingly, the circulating levels of pro-inflammatory cytokines IL-1β, IL-6 and TNF-α in the medium-dose group were significantly increased (*p* = 0.034, *p* < 0.001 and *p* = 0.001, respectively) compared to the CON group (Fig. 1r, s and u). Importantly, the production of pro-inflammatory cytokine TNF-α in the high-dose group was significantly decreased (*p* = 0.008) compared to the medium-dose group (Fig. 1u). In addition, the circulating levels of pro-inflammatory cytokine IL-6 in the high-dose group were significantly increased (*p* = 0.004), and the circulating levels of anti-inflammatory cytokine IL-10 in the low-dose group were significantly decreased (*p* = 0.020), compared to the CON group (Fig. 1s and 1t).

### Dietary GML induces gut microbiota alterations

Three-dimensional principal coordinate analysis (PCoA) plot showed that a significant shift for β-diversity (*p* = 0.024) between CON and GML treatment groups (Fig. 2a). Interestingly, the β-diversity in the medium-dose group was significantly decreased (*p* = 0.033) compared to the CON group, while no significant differences in the CON, low-dose and high-dose groups were observed (Fig. 2b), suggesting that different doses of GML had distinct characteristics to change the communities of gut microbiota. However, no significant differences were observed for α-diversity (Chao 1 index, Observed species, Shannon index and Simpson index, respectively) among all groups (Fig. 2c and Fig. S1).

**Figure 2.**
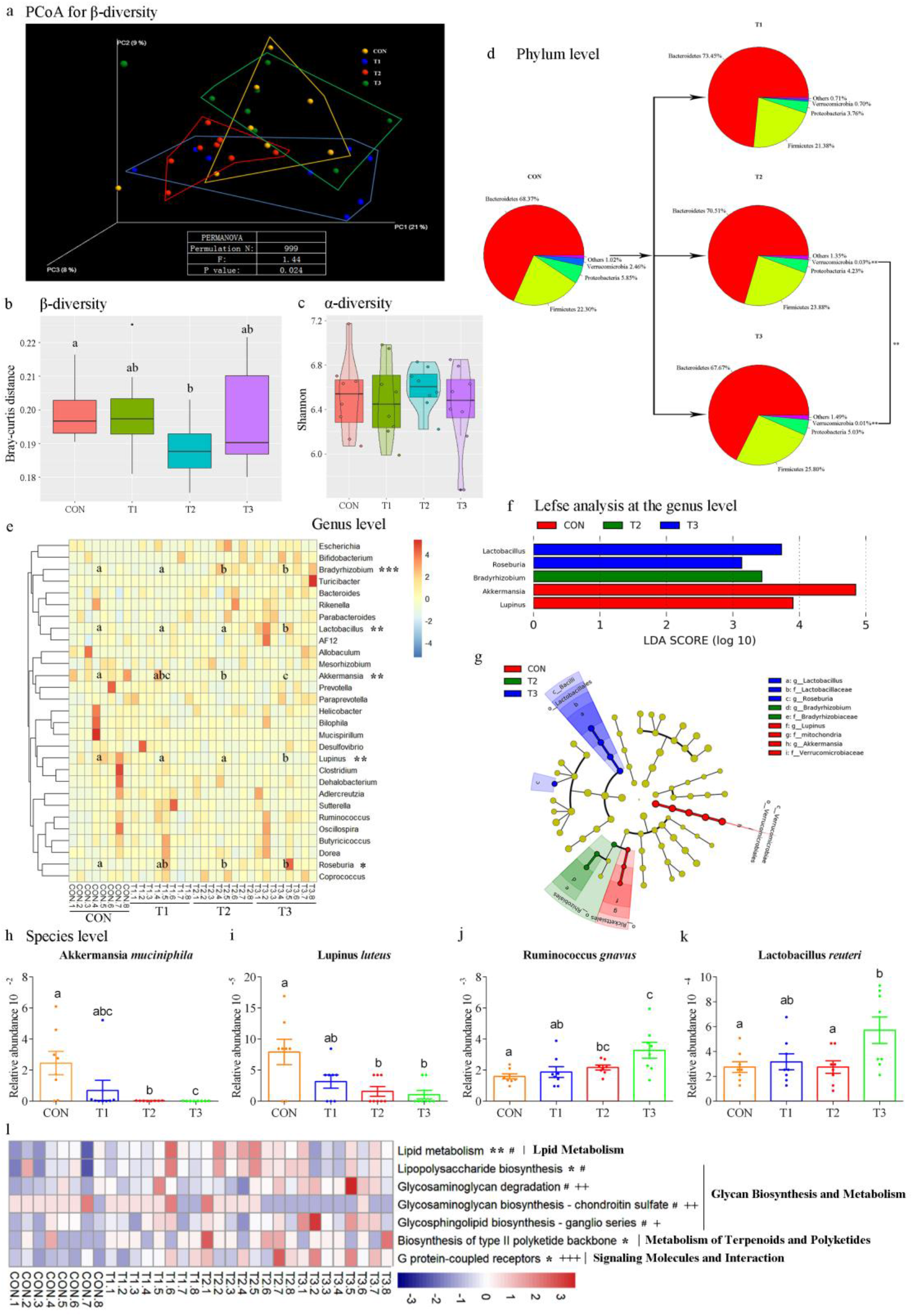
Dietary GML alters gut micobiota diversity, composition and metabolic pathways. **(a)** Three-dimensional principal coordinate analysis (PCoA) plot based on Bray-curtis distances (n = 8/group), **(b)** β-diversity of gut microbiota, **(c)** α-diversity: Shannon index, **(d)** relative abundance of gut microbiota at the phylum level, **(e-g)** Significantly different gut microbiota at the genera level (those with > 0.1% are represented), **(h-k)** Significantly enriched or depleted species and **(l)** the DNA abundances of KEGG pathways in level-3 functional prediction by PICRUSt (n = 8/group). Data are expressed as mean ± SE. Value with asterisk or different alphabet is significantly different (T2 *VS* CON: \**p* < 0.05, \*\**p* < 0.01, \*\*\**p* < 0.001; T3 *VS* T2: ^#^*p* < 0.05, ^##^*p* < 0.01, ^###^*p* < 0.001; T3 VS T1: ^+^p < 0.05, ^++^p < 0.01, ^+++^p < 0.001).

The gut bacterial communities among the CON and GML treatment groups were dominated by two phyla: Bacteroidetes and Firmicutes, and no significant differences were observed (Fig. 2d). However, the relative abundance of the phyla Verrucomicrobia in the medium-dose and high-dose GMLgroup were both significantly decreased (*p* = 0.006 and *p* = 0.006, respectively), compared to the CON group. Moreover, a significant decrease of the phyla Verrucomicrobia in the high-dose GML group were observed compared to the medium-dose group (*p* = 0.008) (Fig. 2d).

Markedly, the genera *Akkermansia* (*p* = 0.006 and *p* = 0.006, respectively) and *Lupinus* (*p* = 0.011 and *p* = 0.006, respectively) in the medium-dose and high-dose groups was significantly decreased, and the genera *Bradyrhizobium* (*p* = 0.013 and *p* = 0.002, respectively) and *Roseburia* (*p* = 0.001 and *p* = 0.042, respectively) in the medium-dose and high-dose groups were significantly increased compared with those of the CON group (Fig. 2e-g). Importantly, the genus *Lactobacillus* in the high-dose group was significantly increased compared with that of the CON, low-dose and medium-dose groups (Fig. 2e-g).

In order to further understand the effects of GML on the modulation of gut microbiota, we identified the significantly enriched or depleted species among CON and GML treatment groups (Fig. 2h-k). Markedly, *Akkermansia muciniphila* in the medium-dose and high-dose groups were significantly decreased by 84.8 times (*p* = 0.006) and 189.1 times (*p* = 0.006), respectively, compared to the CON group (Fig. 2h). *Lupinus luteus* in the medium-dose and high-dose groups were significantly decreased by 4.0 times (*p* = 0.011) and 8.0 times (*p* = 0.006), respectively, compared to the CON group (Fig. 2i). Meanwhile, *Ruminococcus gnavus* in the medium-dose and high-dose GML groups were significantly increased by 1.4 times (*p* = 0.022) and 2.1 times (*p* = 0.007), respectively, compared to that in the CON group (Fig. 2j). Importantly, *Lactobacillus reuteri* in the high-dose group was significantly higher than the medium-dose group (2.1 times, *p* = 0.022) and CON group (2.1 times, *p* = 0.025) (Fig. 2k).

The predictive functions of the microbiota genes were shown in Fig. 2l. The different metabolism pathways belonged to the lipid metabolism, glycan biosynthesis and metabolism, metabolism of terpenoids and polyketides, and signaling molecules and interaction. Interesting, the medium-dose group significantly increased the DNA abundances of 4 predicted metabolism pathways of lipid metabolism (*p* = 0.001), Lipopolysaccharide biosynthesis (*p* = 0.017), biosynthesis of type II polyketide backbone (*p* = 0.031) and G protein-coupled receptors (*p* = 0.022), compared to those in the CON group. Notably, the high-dose group significantly decreased the metabolism pathways of lipid metabolism (*p* = 0.028), Lipopolysaccharide biosynthesis (*p* = 0.027) and glycosaminoglycan biosynthesis-chondroitin sulfate (*p* = 0.042), and significantly increased the metabolism pathways of glycosaminoglycan degradation (*p* = 0.041) and G protein-coupled receptors (*p* = 0.047) compared to those in the medium-dose group.

### Host-microbial interactions driven by GML administration

Co-occurrence network analysis among all groups reveled that *Akkermansia muciniphila* was positively associated with body muscle percentage, and inversely associated with body fat percentage and IL-6. *Lupinus luteus* was positively associated with body muscle percentage, and inversely associated with body fat percentage, IL-6 and TNF-α. *Lactobacillus reuteri* was inversely associated with TG, TC and LDL (Fig. 3a and 3b). In order to further understand the inverse effects of GML on metabolic syndrome, we analysis the host-microbial interactions between medium-dose and high-dose groups (Fig. 3c and 3d). Co-occurrence network analysis between medium-dose and high-dose groups reveled that *Lactobacillus reuteri* was inversely associated with TG, TC, LDL, atherogenic index, LPS, TNF-α and abdominal fat (Fig. 3c and 3d).

**Figure 3.**
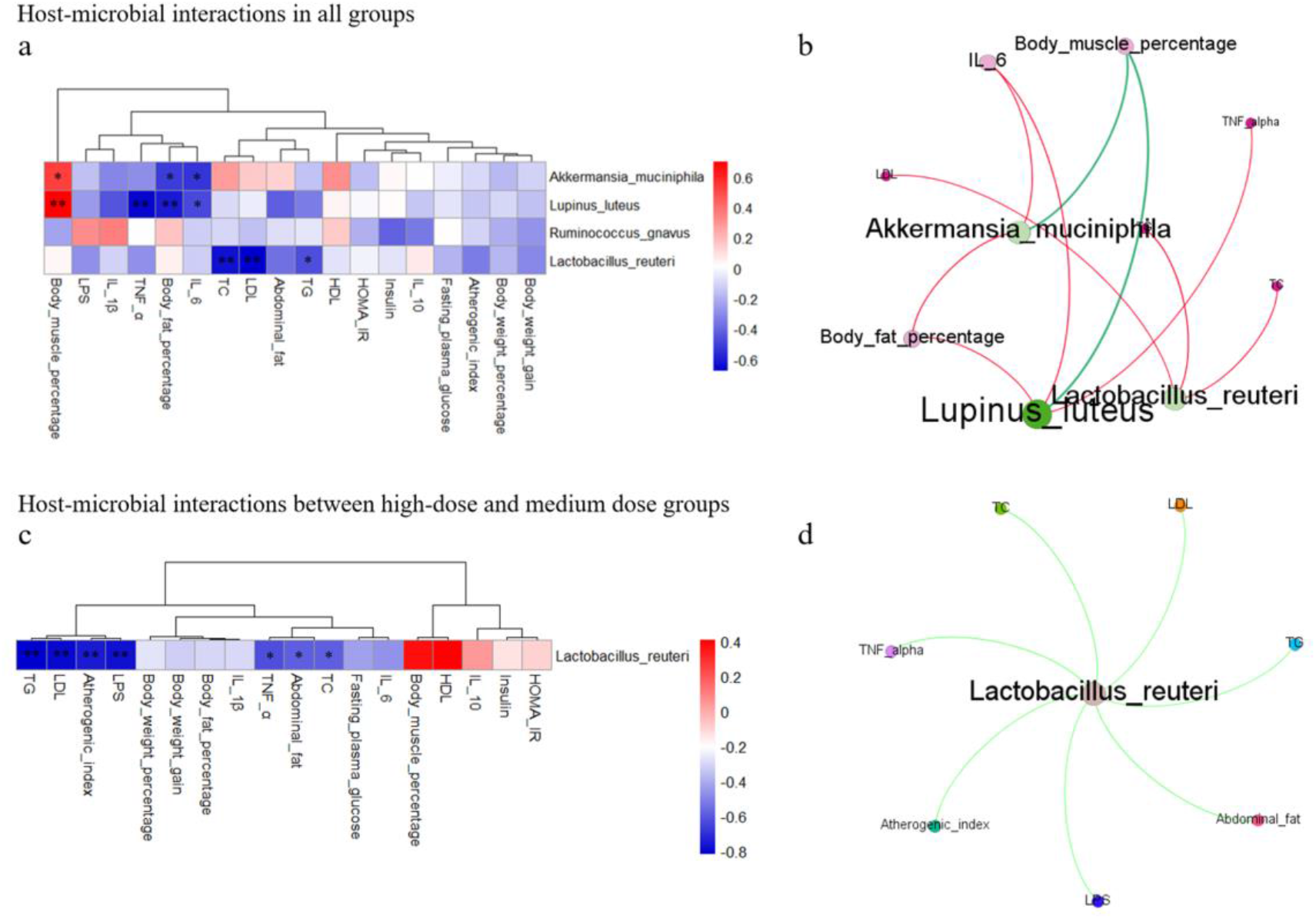
Network analysis revel host-microbial interactions driven by GML administration. Heatmap showing the variation of the association patterns of the significantly enriched or depleted species and host metabolic parameters based on Spearman’s non-parametric rank correlation among all groups **(a)** and between high-dose and medium-dose groups **(c)** (n = 6/group). Host–microbiota interactions networks among all groups **(b)** and between high-dose and medium-dose groups **(d)** were built from Spearman’s non-parametric rank correlation coefficient (red lines indicate positive association, and green lines indicate inverse association). Value with asterisk is significantly different (\**p* < 0.05, \*\**p* < 0.01, \*\*\**p* < 0.001) (n = 6/group). The Benjamini-Hochberg method was used to adjust *p* values for multiple testing.

### Altered gut microbiota is necessary and sufficient for GML-induced distinct effects on metabolic syndrome

Importantly, antibiotics-treated germ-free experiment demonstrated that gut microbiota is necessary for GML-induced metabolic syndrome. Markedly, the body weight (*p* < 0.001), weight gain (*p* < 0.001), body fat percentage ((*p* < 0.001), abdominal fat (*p* < 0.001), TG (*p* < 0.001) and TC (*p* < 0.001) in the antibiotics-treated GML group were significantly decreased compared to those of the 150 mg·kg^−1^ GML group, while no significant differences were observed compared to the CON group (Fig. 4a). In order to further confirm the high-dose GML group induced distinct effect on metabolic syndrome through altered gut microbiota, MetS-mice were built by feeding high-fat diet (HFD) for 8 weeks and *Lactobacillus reuteri* transplantation was conducted for 16 weeks (Fig. 4b). Markedly, the body weight (*p* < 0.001), weight gain (*p* = 0.011), body fat percentage (*p* < 0.001), abdominal fat (*p* < 0.001), TG (*p* < 0.001), TC (*p* = 0.001) and LDL (*p* = 0.002) in the MetS+LR group were significantly decreased compared to those of the MetS group (Fig. 4b).

**Figure 4.**
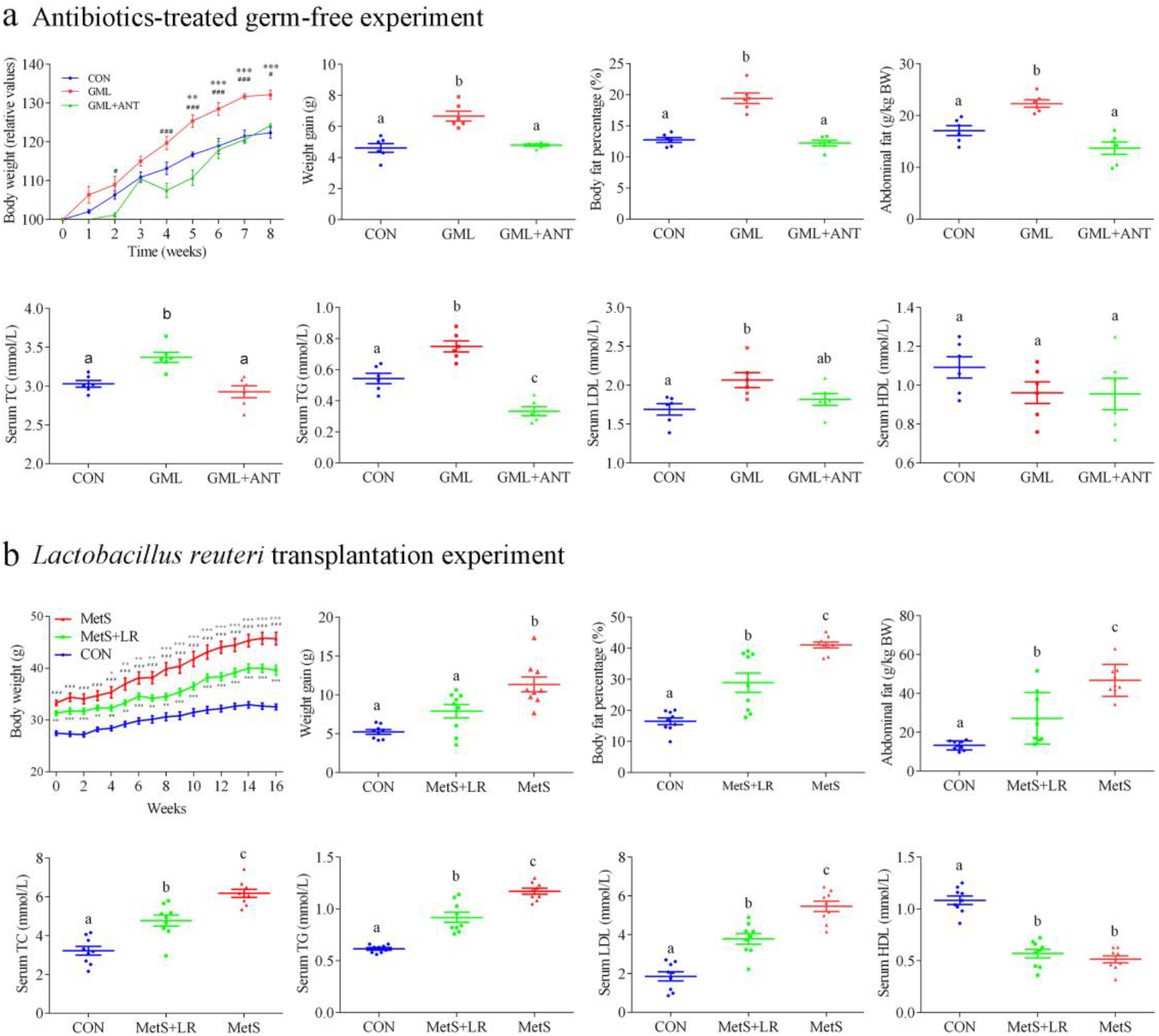
Altered gut microbiota is necessary and sufficient for GML-induced distinct effects on metabolic syndrome. **(a)** Antibiotics-treated germ-free experiment (n = 6/group) and **(b)** *Lactobacillus reuteri* transplantation (n = 9/group). Markers of MetS (including body weight, weight gain, fasting total cholesterol, triglyceride, LDL, HDL and body composition) were analyzed. Data are expressed as mean ± SE. Value with asterisk or different alphabet is significantly different (GML *VS* CON: \**p* < 0.05, \*\**p* < 0.01, \*\*\**p* < 0.001; GML *VS* GML+ANT: ^#^*p* < 0.05, ^##^*p* < 0.01, ^###^*p* < 0.001; MetS+LR *VS* CON: \**p* < 0.05, \*\**p* < 0.01, \*\*\**p* < 0.001; MetS *VS* CON: ^#^*p* < 0.05, ^##^*p* < 0.01, ^###^*p* < 0.001; MetS+LR VS MetS: ^+^*p* < 0.05, ^++^*p* < 0.01, ^+++^*p* < 0.001).

## DISCUSSION

Previous study demonstrated that 150 mg·kg^−1^ food antimicrobial emulsifier-GML consumption induced MetS and gut microbiota dysbiosis in low-fat diet fed mice. However, little is known about the dose-effect of dietary GML modulating the gut microbiome alterations and its impacts on the induction of MetS in a low-fat diet. In the present study, we observed that glycerol monolaurate had no dose-effect on mouse gut microbiome and induced distinct effects on metabolic syndrome via modulation of gut microbiome in low-fat diet, which indicated the importance of reassessment for the safe dosage range of GML in food industry. We propose the new concept that non-specific antibacterial food additives have two-sided effect on gut microbiota contributing to the uncertainties for the incidence of obesity/metabolic syndrome and other chronic inflammatory diseases.

Recently, growing evidence has indicated the relationship between broad use of emulsifying agents and increased societal incidence of metabolic syndrome (18, 19). The NCEP–ATP III proposed that the features of metabolic syndrome were abdominal obesity, raised serum triglycerides and fasting plasma glucose, lower high-density lipoprotein (HDL)-cholesterol and increased blood pressure, and abdominal obesity played a central role in metabolic syndrome. Our previous study demonstrated that 150 mg·kg^−1^ GML consumption induced MetS in low-fat diet fed mice (11). In this study, we also observed that 150 mg·kg^−1^ GML induced mouse MetS in low-fat diet. Meanwhile, 500 mg·kg^−1^ GML significantly alleviated MetS by significantly decreasing the body weight, weight gain, food intake, fat droplet size and percentage of abdominal fat, serum TG, LDL, LPS, TNF-α and atherogenic index, compared to the 150 mg·kg^−1^ GML, indicating that the dietary different doses of GML induced distinct effects on metabolic syndrome without dose-effect in low-fat diet fed mice model.

Emulsifier-induced metabolic syndrome was associated with gut microbiota dysbiosis (6, 8). As already demonstrated in previous studies (6, 20), dietary supplementation with non-caloric artificial sweetener (saccharin) and emulsifiers (CMC and P80), significantly promoted the metabolic syndrome by altering the diversity, composition and function of gut microbiota in mice. However, there are few studies on the dose effect of emulsifiers on gut microbiota. Our study revealed that GML altered the communities of gut microbiota without dose-effect, which is consistent with the changes in physiological phenotypes. It is worth noting that the intestinal mucosal protective genus *Akkermansia* was significantly decreased with dose effect, and the LPS-suppressing genus *Lactobacillus* in the high-dose (500 mg·kg^−1^) GML group were significantly increased respectively compared to the medium-dose (150 mg·kg^−1^) GML group, indicated that high-dose GML alleviated gut microbiota dysbiosis and MetS, at least in part, through significantly increasing LPS-suppressing bacteria (those which can lower the entry of LPS into the circulating system). These findings are consistent with the results of previous studies (21, 22), which demonstrated that increasing LPS-producing bacteria (e.g. *E. coli*), and decreasing LPS-suppressing bacteria (e.g. *Bifidobacterium*) and mucosal protective bacteria (e.g. *Akkermansia*) in gut resulted in metabolic syndrome. Much progress has been made in the identification of beneficial genera, revealing that *Lactobacillus*, *Bifidobacterium*, *Enterococcus faecium*, *Clostridium cluster XIVa*, *Clostridium cluster IV* and *Akkermansia* are mainly gut beneficial genera (23, 24), which have been linked to reducing the intestinal permeability and inflammatory (25), exerting the anti-obesity effects (26) and stimulating the host’s immune system and metabolism (14), and their abundance was correlated significantly with serum LPS levels (27).

Recently, several studies have highlighted that systemic low-grade inflammation resulted from gut dysbiosis, which is now considered as a critical pathological factor underlying many modern chronic diseases, including MetS, type 2 diabetes, cardiovascular disease, cancer and neurodegenerative diseases (28, 29). Systemic low-grade inflammation is characterized by elevated circulating levels of inflammatory cytokines, such as TNF-α, IL-1β, and IL-6 (28) and primarily caused by a gradual increase in plasma endotoxins, particularly LPS. As a consequence, our results showed that high-dose GML moderately reduced systemic low-grade inflammation. Importantly, the serum LPS concentration in high-dose GML group was significantly decreased, compared to the medium-dose GML group, in according with the increase of the LPS-suppressing genus *Lactobacillus* regardless of the relative low level of the mucos-protection genus *Akkermansia* and the relative high level of the LPS-producing genus *Bradyrhizobium*, indicating that the serum LPS is an important link in the cross-talk between the gut microbiota and the host inflammation, and the supplementation of high-dose GML might reduce gut LPS-related microbiota dysbiosis to decrease in the serum LPS concentration and further alleviate systemic low-grade inflammation and MetS. Thus, a better understanding of the dose-effects of dietary GML for modulating gut microbiota would be important for evaluating the safety of its use in consumed foods.

Specific and significant increase of Akkermansia *muciniphila* (widely considered as mucos-protection bacterium) in the gut was important for reducing the host susceptibility to metabolic syndrome and systemic low-grade inflammation (30, 31), but our results suggested that GML treatments could significantly decrease Akkermansia *muciniphila* and Lupinus *luteus*, and increase *Ruminococcus gnavus,* which were adverse to reduce the serum LPS concentration and pro-inflammatory cytokines (32), and might result in MetS, obesity and metabolic endotoxemia (33). Notably, high-dose GML significantly increased *Lactobacillus reuteri* compared to the medium-dose GML. *Lactobacillus reuteri* has recently been shown to have an important role in the prevention against obesity (34), systemic low-grade inflammation (35) and insulin resistance in mice (36), the improvement of MetS and the production of aryl hydrocarbon receptor ligand in the intestine of mice (37). This production plays a vital role in the improvement of intestinal barrier function and secretion of the incretin hormone Glucagon-Like Peptide (GLP)-1 (37). Accordantly, co-occurrence network analysis in our study revealed that *Lactobacillus reuteri* was positively associated with the metabolic improvement driven by high-dose GML administration. PICRUSt function prediction further indicated that the gut microbiota in the high-dose GML group had a significantly lower capacity of the lipid metabolism and lipopolysaccharide biosynthesis compared to those in the medium-dose GML group, which would decrease host energy harvest and circulating inflammation level contributing to the prevention of obesity/metabolic syndrome and other chronic inflammatory diseases (6).

In order to further understand whether GML-mediated gut microbiota alterations were necessary for the induction of MetS, we conducted the antibiotics-treated germ-free experiment and *Lactobacillus reuteri* transplantation. Notably, antibiotics-treated germ-free experiment demonstrated that gut microbiota was necessary for GML-induced metabolic syndrome. Further, *Lactobacillus reuteri* transplantation demonstrated that *Lactobacillus reuteri* significantly ameliorated high fat diet-induced MetS, indicating that altered gut microbiota was necessary and sufficient for GML-induced distinct effects on metabolic syndrome. In summary, these findings indicated that different doses of GML in low fat diet impacted gut microbiome inducing distinct effects on metabolic syndrome.

In conclusion, the present study indicates that dietary different doses of GML in low fat diet had the potential to induce distinct effects on metabolic syndrome via modulation of gut microbiota. Our study proposes the new concept that non-specific antibacterial food additives have two-sided effect on gut microbiota contributing to the uncertainties for the incidence of obesity/metabolic syndrome and other chronic inflammatory diseases, thereby calling for reassessing the safe dosage of GML and other non-specific antibacterial food additives. However, the underlying mechanism of the distinct effects on metabolic syndrome by GML needs to be further explored.

## MATERIALS AND METHODS

### Ethics statement

Four weeks-old female C57BL/6 mice were purchased from SLAC Laboratory Animal (Shanghai, China) and housed in the Zhejiang University animal facility in a specific pathogen free level room in hard top cages with two or three mice per cage. Mice were maintained at 22–24 °C in a 12-h light/dark cycle and allowed *ad libitum* access to food and water. All procedures were conducted in accordance with institutional guidelines and were approved by the Animal Ethical Committee of Zhejiang University (reference protocol number ZJU-BEFS-2015012) and Ethics Committee of Westlake University (reference protocol number 18-003-ZJS).

### GML treatment

Mice were randomly allocated into four groups (n = 8 per group): (1) the basal Purina rodent chow diet no. P1100F (SLACOM Inc, Shanghai) as the control group (CON); (2) the basal diet supplemented with 50 mg·kg^−1^ GML as the low-dose group (T1 group); (3) the basal diet supplemented with 150 mg·kg^−1^ GML as the median-dose group (T2 group); (4) the basal diet supplemented with 500 mg·kg^−1^ GML as the high-dose group (T3 group) (Table S1). Each treatment has 3 cages (3 or 2 mice per cage). Body weights were individually measured every week and expressed as a percentage compared to the initial body weight (0 day) defined as 100%. Fresh faeces were collected every week for subsequent analysis. After 8 weeks of GML treatment, mice were fasted for 12 h, and blood was collected by retrobulbar intraorbital capillary plexus. Mice were then euthanized by i.p. injection of pentobarbital (200 mg·kg^−1^), and abdominal adipose weight were measured. Organs were collected for subsequent analysis. Incorporation of GML into the basic diet (Purina Rodent Chow diet no. P1100F) and maintaining total energy balance between diets by replacing isoenergetic fat with GML were performed by SLACOM Inc. (Shanghai, China).

### DNA extraction and 16S rRNA gene sequencing

DNA extractions from the 8-week fecal samples (n = 8 per group) were performed using the Qiagen QIAmp DNA stool extraction kit (Qiagen) following the manufacturer’s protocol. PCR amplification of 16S rRNA V4 regions were conducted by using the 515F/806R bacterial primer with the barcode (515F: 5’-GTGCCAGCMGCCGCGGTAA-3’ and 806R: 5’-GGAC TACHVGGGTWTCTAAT-3’) (38). PCR products were mixed in equidensity ratios. Sequencing libraries were generated using NEB Next® Ultra™ DNA Library Prep Kit for Illumina (NEB, USA) following manufacturer’s recommendations and index codes were added. The library quality was assessed on the Qubit@ 2.0 Fluorometer (Thermo Scientific) and Agilent Bioanalyzer 2100 system. At last, the library was sequenced on an Illumina MiSeq platform and 250 bp paired-end reads were generated (11). Details of the 16S rRNA sequence preparation and analysis (39) are available in Text 1 in supplemental materials.

### Serum lipid parameters and fasting blood glucose testing

Total serum cholesterol (TC), triglycerides (TG), high-density lipoprotein cholesterol (HDL-C), low-density lipoprotein cholesterol (LDL-C), fasting plasma glucose (FPG) and insulin (FINS) were determined using a kit from Nanjin Jiangcheng Bioengineering Institute (Jiangshu, China), following the manufacturer’s instructions. Atherogenic index = (TC− HDL-C)/HDL-C (40).

### Body composition analysis

Body fat percentage were determined with the nuclear magnetic resonance system using a Body Composition Analyzer MesoQMR06-100H (Suzhou Niumag Analytical Instrument Corporation, China), as described previously (11).

### Abdominal adipose tissue histopathology

Mouse abdominal adipose tissue samples (n=5) were fixed in 10% buffered formalin for 24 h at room temperature and then embedded in paraffin, respectively. Tissues (five sections per mouse) were sectioned at 5 μm thickness and stained with haematoxylin and eosin (H&E) using standard protocols, and the number and size of stained fat droplets in abdominal adipose tissue were analysed by the Image J software (National Institutes of Health, USA) (41).

### Measurement of systemic cytokine levels and serum LPS concentration

Enzyme-linked immunosorbent assay (ELISA) kits were used to determine serum levels of IL-1β, IL-6, TNF-α and IL-10 (eBioscience), according to the manufacturers’ instructions. Serum LPS concentrations were measured with a ToxinSensor Chromogenic Limulus Amebocyte Lysate (LAL) Endotoxin Assay Kit (GenScript, Piscataway, NJ), following the manufacturer’s instructions. Before measuring LPS concentrations, the serum was heated for 10 min at 70 °C to minimize inhibition or enhancement by contaminating proteins. LAL reagents were added to serum and incubated at 37 °C for 45 min, and the absorbance was read at 545 nm. All samples were validated for recovery and internal coefficient variation using known amounts of LPS.

### Antibiotics-treated germ-free experiments

Mice were randomly divided into three groups (n = 6 per group) : (1) the basal Purina rodent chow diet as the control group (CON); (2) the basal diet supplemented with 150 mg·kg^−1^ GML as the GML treatment group (GML); (3) the basal diet supplemented with 150 mg·kg^−1^ GML and a broad spectrum mixed antibiotics containing ampicillin (1 g/L), vancomycin (500 mg/L), neomycin sulfate (1 g/L) and metronidazole (100 mg/kg) (added to drinking water) as the antibiotics-treated germ-free group (GML+ANT). After 8 weeks experiments, mice with or without antibiotics treatment were subjected to analysis for markers of MetS (including fasting total cholesterol, triglyceride, LDL, HDL and body composition).

### Gut microbiota transplantation

*Lactobacillus reuteri* was purchased from the Chinese academy of agricultural sciences and identified as *Lactobacillus reuteri* ZLR003 by whole 16S rDNA sequencing. *L. reuteri* was cultured on Man–Rogosa–Sharpe (MRS) medium, and subjected to a serious of centrifugation steps to separate the MRS cultured broth into *L. reuteri*-riched supernatant (0.9% saline solution) as described previously (42). MetS-mice were built by feeding high-fat diet (HFD) for 8 weeks. Mice (n = 9) were fed with the basal chow diet as the normal group (CON). Validation of successful establishment of MetS-mice was performed as described previously (43). MetS-mice were randomly divided into two groups (n = 9 per group): (1) the HFD group (MetS); (2) the HFD supplemented with *L. reuteri* (MetS+LR). For the transplantation with *L. reuteri*, MetS-mice were gavaged daily (100 μl 0.9% saline solution) with 10^10^ CFU/ml of *L. reuteri* or vehicle (0.9% saline solution) for 7 days and then orally administered by drinking water (10^9^ CFU/ml or normal water respectively) for 16 weeks. After treatment for 16 weeks, mice were subjected to analysis for markers of MetS (including fasting total cholesterol, triglyceride, LDL, HDL and body composition).

### Statistical analysis

The data of results were collected individually, calculated for each treatment, and expressed as mean ± standard error (SE). Statistical differences between more than two groups were evaluated by one-way analysis of variance (ANOVA) with Tukey’s multiple comparison post-tests. Data between two treatments were determined using the unpaired two-tailed T-test in GraphPad Prism version 6 (GraphPad Software, La Jolla, CA). Principal coordinate analysis (PCoA) based on Bray-Curtis distance and permutational multivariate analysis of variance (PERMANOVA) were performed to compare the dissimilarity of global microbiota composition. Microbiome-based biomarker discovery was performed with LEfSe using the online galaxy server (https://huttenhower.sph.harvard.edu/galaxy/). LDA scores derived from the LEfSe analysis were used to show the relationship between taxa using a cladogram (circular hierarchical tree) of significantly increased or decreased bacterial taxa in the gut microbiota between groups. Levels of the cladogram from the inner to outer rings: phylum, class, order, family, and genus. Color codes indicate the groups, and letters indicate the taxa that contribute to the uniqueness of the corresponding groups at an LDA of > 2.0. Data from high-throughput sequence were analyzed in QIIME-2-2020.2 and R 3.6.3. The co-occurrence network analysis based on the spearman correlation to demonstrate the host-microbial interactions driven by GML adminstration. The Benjamini-Hochberg method was used to adjust *p* values for multiple testing. The networks were further visualized in Gephi software version 0.9.2. Data were checked for heterogeneous variance with the Brown-Forsythe tests or the F test. *p*-value < 0.05 was considered significant and 0.05 < *p*-value ≤ 0.10 were discussed as tendencies.

## Availability of data and materials

The raw data for 16 S rRNA gene sequences are available in the CNSA (https://db.cngb.org/cnsa/) of CNGBdb at accession number CNS0267006.

## ACKNOWLEDGEMENTS

We would like to thank Suzhou Niumag Analytical Instrument Corporation for mouse body composition analysis. The present work was supported by the Natural Science Foundation of Zhejiang Province (Grant No. LD19C200001 and LQ19C200005), National Natural Science Foundation of China (Grant No. 81903316), Zhejiang Ten-thousand Talents Program (Grant No. 101396522001), Technology and Achievement Transformation Project of Hangzhou, China (Grant No. 20161631E01), Zhejiang University New Rural Development Research Institute Agricultural Technology Promotion Fund (Grant No. 2017006) and Yunnan Provincial Science and Technology Department-Applied Basic Research Joint Special Funds of Yunnan University of Traditional Chinese Medicine (Grant No. 2017FF116(−001)).

## References

1. Eckel RH, Grundy SM, Zimmet PZ. 2005. The metabolic syndrome. Lancet 365:1415–28.

2. Cornier MA, Dabelea D, Hernandez TL, Lindstrom RC, Steig AJ, Stob NR, Van Pelt RE, Wang H, Eckel RH. 2008. The metabolic syndrome. Endocr Rev 29:777–822.

3. Despres JP, Lemieux I. 2006. Abdominal obesity and metabolic syndrome. Nature 444:881–7.

4. Aguilar M, Bhuket T, Torres S, Liu B, Wong RJ. 2015. Prevalence of the metabolic syndrome in the United States, 2003-2012. JAMA 313:1973–4.

5. Athyros VG, Ganotakis ES, Elisaf M, Mikhailidis DP. 2005. The prevalence of the metabolic syndrome using the National Cholesterol Educational Program and International Diabetes Federation definitions. Current Medical Research and Opinion 21:1157–1159.

6. Chassaing B, Koren O, Goodrich JK, Poole AC, Srinivasan S, Ley RE, Gewirtz AT. 2015. Dietary emulsifiers impact the mouse gut microbiota promoting colitis and metabolic syndrome. Nature 519:92–6.

7. Chassaing B, Van de Wiele T, De Bodt J, Marzorati M, Gewirtz AT. 2017. Dietary emulsifiers directly alter human microbiota composition and gene expression ex vivo potentiating intestinal inflammation. Gut 66:1414–1427.

8. Viennois E, Merlin D, Gewirtz AT, Chassaing B. 2017. Dietary Emulsifier-Induced Low-Grade Inflammation Promotes Colon Carcinogenesis. Cancer Research 77:27–40.

9. Kabara JJ, Swieczkowski DM, Conley AJ, Truant JP. 1972. Fatty acids and derivatives as antimicrobial agents. Antimicrob Agents Chemother 2:23–8.

10. Zhang H, Wei H, Cui Y, Zhao G, Feng F. 2009. Antibacterial interactions of monolaurin with commonly used antimicrobials and food components. J Food Sci 74:M418–21.

11. Jiang Z, Zhao M, Zhang H, Li Y, Liu M, Feng F. 2018. Antimicrobial Emulsifier-Glycerol Monolaurate Induces Metabolic Syndrome, Gut Microbiota Dysbiosis, and Systemic Low-Grade Inflammation in Low-Fat Diet Fed Mice. Mol Nutr Food Res 62.

12. Zhao MJ, Cai HY, Jiang ZL, Li Y, Zhong H, Zhang H, Feng FQ. 2019. Glycerol-Monolaurate-Mediated Attenuation of Metabolic Syndrome is Associated with the Modulation of Gut Microbiota in High-Fat-Diet-Fed Mice. Molecular Nutrition & Food Research 63.

13. Clemente JC, Ursell LK, Parfrey LW, Knight R. 2012. The impact of the gut microbiota on human health: an integrative view. Cell 148:1258–70.

14. Qin J, Li Y, Cai Z, Li S, Zhu J, Zhang F, Liang S, Zhang W, Guan Y, Shen D, Peng Y, Zhang D, Jie Z, Wu W, Qin Y, Xue W, Li J, Han L, Lu D, Wu P, Dai Y, Sun X, Li Z, Tang A, Zhong S, Li X, Chen W, Xu R, Wang M, Feng Q, Gong M, Yu J, Zhang Y, Zhang M, Hansen T, Sanchez G, Raes J, Falony G, Okuda S, Almeida M, LeChatelier E, Renault P, Pons N, Batto JM, Zhang Z, Chen H, Yang R, Zheng W, Li S, Yang H, et al. 2012. A metagenome-wide association study of gut microbiota in type 2 diabetes. Nature 490:55–60.

15. Vrieze A, Van Nood E, Holleman F, Salojarvi J, Kootte RS, Bartelsman JF, Dallinga-Thie GM, Ackermans MT, Serlie MJ, Oozeer R, Derrien M, Druesne A, Van Hylckama Vlieg JE, Bloks VW, Groen AK, Heilig HG, Zoetendal EG, Stroes ES, de Vos WM, Hoekstra JB, Nieuwdorp M. 2012. Transfer of intestinal microbiota from lean donors increases insulin sensitivity in individuals with metabolic syndrome. Gastroenterology 143:913–6 e7.

16. Burrello C, Giuffre MR, Macandog AD, Diaz-Basabe A, Cribiu FM, Lopez G, Borgo F, Nezi L, Caprioli F, Vecchi M, Facciotti F. 2019. Fecal Microbiota Transplantation Controls Murine Chronic Intestinal Inflammation by Modulating Immune Cell Functions and Gut Microbiota Composition. Cells 8.

17. Kaliannan K, Li XY, Wang B, Pan Q, Chen CY, Hao L, Xie SF, Kang JX. 2019. Multi-omic analysis in transgenic mice implicates omega-6/omega-3 fatty acid imbalance as a risk factor for chronic disease. Communications Biology 2.

18. Roca-Saavedra P, Mendez-Vilabrille V, Miranda JM, Nebot C, Cardelle-Cobas A, Franco CM, Cepeda A. 2018. Food additives, contaminants and other minor components: effects on human gut microbiota-a review. Journal of Physiology and Biochemistry 74:69–83.

19. Partridge D, Lloyd KA, Rhodes JM, Walker AW, Johnstone AM, Campbell BJ. 2019. Food additives: Assessing the impact of exposure to permitted emulsifiers on bowel and metabolic health-introducing the FADiets study. Nutr Bull 44:329–349.

20. Suez J, Korem T, Zeevi D, Zilberman-Schapira G, Thaiss CA, Maza O, Israeli D, Zmora N, Gilad S, Weinberger A, Kuperman Y, Harmelin A, Kolodkin-Gal I, Shapiro H, Halpern Z, Segal E, Elinav E. 2014. Artificial sweeteners induce glucose intolerance by altering the gut microbiota. Nature 514:181–6.

21. Kaliannan K, Wang B, Li XY, Kim KJ, Kang JX. 2015. A host-microbiome interaction mediates the opposing effects of omega-6 and omega-3 fatty acids on metabolic endotoxemia. Sci Rep 5:11276.

22. Zhao SQ, Liu W, Wang JQ, Shi J, Sun YK, Wang WQ, Ning G, Liu RX, Hong J. 2017. Akkermansia muciniphila improves metabolic profiles by reducing inflammation in chow diet-fed mice. Journal of Molecular Endocrinology 58:1–14.

23. Louis P, Flint HJ. 2009. Diversity, metabolism and microbial ecology of butyrate-producing bacteria from the human large intestine. Fems Microbiology Letters 294:1–8.

24. Louis P, Young P, Holtrop G, Flint HJ. 2010. Diversity of human colonic butyrate-producing bacteria revealed by analysis of the butyryl-CoA:acetate CoA-transferase gene. Environmental Microbiology 12:304–314.

25. Martin R, Chain F, Miquel S, Lu J, Gratadoux JJ, Sokol H, Verdu EF, Bercik P, Bermudez-Humaran LG, Langella P. 2014. The Commensal Bacterium Faecalibacterium prausnitzii Is Protective in DNBS-induced Chronic Moderate and Severe Colitis Models. Inflammatory Bowel Diseases 20:417–430.

26. Clemente JC, Ursell LK, Parfrey LW, Knight R. 2012. The Impact of the Gut Microbiota on Human Health: An Integrative View. Cell 148:1258–1270.

27. Duncan SH, Belenguer A, Holtrop G, Johnstone AM, Flint HJ, Lobley GE. 2007. Reduced dietary intake of carbohydrates by obese subjects results in decreased concentrations of butyrate and butyrate-producing bacteria in feces. Applied and Environmental Microbiology 73:1073–1078.

28. Hotamisligil GS. 2006. Inflammation and metabolic disorders. Nature 444:860–7.

29. Ruiz-Nunez B, Pruimboom L, Dijck-Brouwer DA, Muskiet FA. 2013. Lifestyle and nutritional imbalances associated with Western diseases: causes and consequences of chronic systemic low-grade inflammation in an evolutionary context. J Nutr Biochem 24:1183–201.

30. Depommier C, Everard A, Druart C, Plovier H, Van Hul M, Vieira-Silva S, Falony G, Raes J, Maiter D, Delzenne NM, de Barsy M, Loumaye A, Hermans MP, Thissen JP, de Vos WM, Cani PD. 2019. Supplementation with Akkermansia muciniphila in overweight and obese human volunteers: a proof-of-concept exploratory study. Nature Medicine 25:1096–+.

31. Mo Q, Fu A, Deng L, Zhao M, Li Y, Zhang H, Feng F. 2019. High-dose Glycerol Monolaurate Up-Regulated Beneficial Indigenous Microbiota without Inducing Metabolic Dysfunction and Systemic Inflammation: New Insights into Its Antimicrobial Potential. Nutrients 11.

32. Dao MC, Everard A, Aron-Wisnewsky J, Sokolovska N, Prifti E, Verger EO, Kayser BD, Levenez F, Chilloux J, Hoyles L, Dumas ME, Rizkalla SW, Dore J, Cani PD, Clement K, Consortium M-O. 2016. Akkermansia muciniphila and improved metabolic health during a dietary intervention in obesity: relationship with gut microbiome richness and ecology. Gut 65:426–436.

33. Li J, Lin SQ, Vanhoutte PM, Woo CW, Xu AM. 2016. Akkermansia Muciniphila Protects Against Atherosclerosis by Preventing Metabolic Endotoxemia-Induced Inflammation in Apoe(-/-) Mice. Circulation 133:2434–+.

34. Chen LH, Chen YH, Cheng KC, Chien TY, Chan CH, Tsao SP, Huang HY. 2018. Antiobesity effect of Lactobacillus reuteri 263 associated with energy metabolism remodeling of white adipose tissue in high-energy-diet-fed rats. Journal of Nutritional Biochemistry 54:87–94.

35. Tenorio-Jimenez C, Martinez-Ramirez MJ, Del Castillo-Codes I, Arraiza-Irigoyen C, Tercero-Lozano M, Camacho J, Chueca N, Garcia F, Olza J, Plaza-Diaz J, Fontana L, Olivares M, Gil A, Gomez-Llorente C. 2019. Lactobacillus reuteri V3401 Reduces Inflammatory Biomarkers and Modifies the Gastrointestinal Microbiome in Adults with Metabolic Syndrome: The PROSIR Study. Nutrients 11.

36. Hsieh FC, Lee CL, Chai CY, Chen WT, Lu YC, Wu CS. 2013. Oral administration of Lactobacillus reuteri GMNL-263 improves insulin resistance and ameliorates hepatic steatosis in high fructose-fed rats. Nutrition & Metabolism 10.

37. Natividad JM, Agus A, Planchais J, Lamas B, Jarry AC, Martin R, Michel ML, Chong-Nguyen C, Roussel R, Straube M, Jegou S, McQuitty C, Le Gall M, da Costa G, Lecornet E, Michaudel C, Modoux M, Glodt J, Bridonneau C, Sovran B, Dupraz L, Bado A, Richard ML, Langella P, Hansel B, Launay JM, Xavier RJ, Duboc H, Sokol H. 2018. Impaired Aryl Hydrocarbon Receptor Ligand Production by the Gut Microbiota Is a Key Factor in Metabolic Syndrome. Cell Metabolism 28:737–+.

38. Martinezmurcia AJ, Acinas SG, Rodriguezvalera F. 1995. Evaluation of Prokaryotic Diversity by Restrictase Digestion of 16s Rdna Directly Amplified from Hypersaline Environments. Fems Microbiology Ecology 17:247–255.

39. Bolyen E, Rideout JR, Dillon MR, Bokulich N, Abnet CC, Al-Ghalith GA, Alexander H, Alm EJ, Arumugam M, Asnicar F, Bai Y, Bisanz JE, Bittinger K, Brejnrod A, Brislawn CJ, Brown CT, Callahan BJ, Caraballo-Rodriguez AM, Chase J, Cope EK, Da Silva R, Diener C, Dorrestein PC, Douglas GM, Durall DM, Duvallet C, Edwardson CF, Ernst M, Estaki M, Fouquier J, Gauglitz JM, Gibbons SM, Gibson DL, Gonzalez A, Gorlick K, Guo JR, Hillmann B, Holmes S, Holste H, Huttenhower C, Huttley GA, Janssen S, Jarmusch AK, Jiang LJ, Kaehler BD, Bin Kang K, Keefe CR, Keim P, Kelley ST, Knights D, et al. 2019. Reproducible, interactive, scalable and extensible microbiome data science using QIIME 2. Nature Biotechnology 37:852–857.

40. Mohammadi A, Oshaghi EA. 2014. Effect of garlic on lipid profile and expression of LXR alpha in intestine and liver of hypercholesterolemic mice. J Diabetes Metab Disord 13:20.

41. Chang CJ, Lin CS, Lu CC, Martel J, Ko YF, Ojcius DM, Tseng SF, Wu TR, Chen YYM, Young JD, Lai HC. 2015. Ganoderma lucidum reduces obesity in mice by modulating the composition of the gut microbiota. Nature Communications 6.

42. Hemarajata P, Gao C, Pflughoeft KJ, Thomas CM, Saulnier DM, Spinler JK, Versalovic J. 2013. Lactobacillus reuteri-Specific Immunoregulatory Gene rsiR Modulates Histamine Production and Immunomodulation by Lactobacillus reuteri. Journal of Bacteriology 195:5567–5576.

43. Fraulob JC, Ogg-Diamantino R, Fernandes-Santos C, Aguila MB, Mandarim-de-Lacerda CA. 2010. A Mouse Model of Metabolic Syndrome: Insulin Resistance, Fatty Liver and Non-Alcoholic Fatty Pancreas Disease (NAFPD) in C57BL/6 Mice Fed a High Fat Diet. J Clin Biochem Nutr 46:212–23.

